# Neuromodulatory influences on propagation of brain waves along the unimodal-transmodal gradient

**DOI:** 10.1101/2024.10.06.616793

**Authors:** Veronica Maki-Marttunen, Sander Nieuwenhuis

## Abstract

Brain activity fluctuates over time, and understanding the factors that influence such fluctuations is important to understand the flexible nature of the brain and cognition. Growing evidence suggests that fMRI brain activity shows spatio-temporal patterns of propagation following specific gradients. In particular, activity around global peaks propagates as a travelling wave following a gradient from unimodal to associative areas. Some properties of these travelling waves seem to be related to behavioral and arousal states, however their meaning remains unknown. Here we assess the possibility that travelling waves explain the finding that there are specific time points when the brain presents larger brain integration. We reasoned that a faster speed of propagation would be related to more brain integration as measured with fMRI. Furthermore, we explored whether increased pupil-linked arousal, which has been related to more integration in specific brain regions, would be increased during periods of whole brain propagation. To test these hypotheses, we detected brain travelling waves and characterized them in terms of speed, directionality and ratio. We compared these features between different task conditions, and after a pharmacological challenge affecting neuromodulatory tone. We then studied the relation between travelling wave speed, pupil size and a graph-based measure of brain integration. Our results suggest that neuromodulatory tone affects travelling wave propagation, and that this propagation reflects changes in arousal and integrated functional connectivity features. This study provides a novel view of brain dynamics in terms of the effects of neruomodulatory influences across time scales.

## Introduction

A still burning question in neuroscience is how the brain maintains high flexibility and adaptability to deal with external and internal demands despite a relatively fixed anatomical structure (Yin and Kaiser, 2021). To answer this question, the field of brain dynamics has grown significatively in the past decade (Lurie et al. 2020, Iraji et al. 2021, Keilholz et al. 2017, Hutchison et al. 2013, Calhoun et al. 2014, Cabral et al. 2014, Engel and Steinmetz 2019). Studies using functional magnetic resonance imaging (fMRI) initially focused on measures of functional connectivity, showing how correlations in blood oxygen level-dependent (BOLD) activity across multiple brain regions change over time (Allen et al. 2014, Chang and Glover 2010, see Preti et al. 2017 for a review). One key aspect that emerged from that work was that, over the course of an experiment, brain activity fluctuates between periods of network-level integration, a state in which there are relatively strong interconnections between network modules, and periods of network-level segregation, a state in which there is a relatively high network modularity (Betzel et al. 2016, Keilholz et al. 2013, Shine and Poldrack 2018). These dynamics have been associated with the flexible nature of mental experience (MacDowell et al. 2022), but how states of integration and segregation correspond to neural events in global brain activity is still unclear.

In the past years, research on whole-brain dynamics in animals and humans has begun to directly examine fluctuations in the BOLD signal (in rodents: Liu and Zhang 2019, Mitra et al. 2018, Vafaii et al. 2024, MacDowell and Buschman 2020; in humans: see Meyer-Baese et al. 2022 for a review). Growing evidence using a variety of approaches suggests the presence of propagating waves at a macroscopic scale and in the slow frequencies (Matsui et al. 2016, Raut et al. 2020, Yousefi and Keilholz 2021), which appear to be relevant for cognition (Xu et al. 2023). In particular, a number of fMRI and electrophysiology studies in rodents, primates and humans suggests that activity seems to propagate through the brain as travelling waves following a gradient from unimodal to associative areas (Matsui et al. 2016, Gu et al. 2021). An interesting possibility has thus been suggested that the propagation of activity across this gradient could give rise to some of the hallmark findings in studies of functional connectivity (Shahsavarani et al. 2023, Raut et al. 2023). The canonical approach of identifying correlated “networks” of activity may be influenced by the position of these networks along what is actually a continuous gradient underlying observed travelling waves. If this is the case, one prediction would be that a change in the *speed of propagation* must cause variations in the correlation between networks, which would be reflected in changes in overall integration (Mäki-Marttunen 2023). That is, if activity propagates faster, it would result in more temporal alignment between regions’ time courses, manifested as higher integration. Finding such evidence would lend further support for the slow propagation of activity as a comprehensive framework to study brain dynamics and for our understanding of the factors that make the brain highly flexible.

What can cause changes in the propagation speed of travelling waves? One possible factor is the level of neural excitability as set by neuromodulatory systems (Raut et al. 2023). Neuromodulatory systems such as the dopamine system and noradrenaline system work at fast and slow time scales and influence large parts of the brains through their wide-spread projections (McCormick et al. 2020, van den Brink et al. 2019). It is well known that low levels of neuromodulatory tone are related to lower excitability and more neural synchronization, while intermediate levels are related to higher excitability and responsivity, and an overall desynchronization (Aston-Jones and Cohen 2005). A computational study has assessed the effect of excitability on travelling waves, and they found that larger excitability was associated with faster propagation (Bhattacharya et al. 2021). Studies in humans offer indirect evidence, by showing that increases or decreases in pupil size during the course of an experiment are associated with more or less brain integration, respectively (Shine 2019, Shine et al. 2016, Mäki-Marttunen 2021). However, a relation between neuromodulation, global brain activity dynamics and measures of integration remains to be investigated in the context of macroscopic travelling waves in the human brain.

In the current study, we aimed to test these hypotheses in a concurrent fMRI-pupillometry study including a pharmacological challenge of neuromodulatory systems. We first detected travelling waves and characterized them in terms of speed, directionality and ratio of top-down to bottom-up propagations. We compared these features between resting state and task performance, and after placebo or atomoxetine manipulation. Atomoxetine is a blocker of the noradrenaline transporter and is associated with an increase in the free levels of catecholamines (i.e. dopamine and noradrenaline, (Bymaster et al., 2002; Swanson et al., 2006; Koda et al., 2010). We then studied the relation between travelling waves, pupil size (a measure of neuromodulatory tone; Joshi and Gold, 2020, Lloyd et al., 2023) and the participation coefficient, a graph-based measure of brain integration that is sensitive to fluctuations in neuromodulatory inputs (Shine et al. 2016, 2018, Mäki-Marttunen 2021). Based on the predictions laid out in Mäki-Marttunen 2023, we hypothesized that (i) pharmacollogically increased levels of catecholamines would be associated with faster travelling waves, that (ii) faster travelling wave speed would be associated with a more integrated network state, and that (iii) the presence of travelling waves would be associated with larger pupil size. Our results provide the first evidence in humans that neuromodulation is related to global brain activity propagation and that these events of propagation can explain why the level of integration between functional networks fluctuates over time. This study is an important step towards understanding how neuromodulatory systems endow the brain with a large flexibility by modulating patterns of global communication.

## Methods

### Participants

Thirty-six individuals (average age: 23 years old, range: 18–29 years, 14 male) were recruited and medically screened by a physician for physical health and drug contraindications. Exclusion criteria included: standard contraindications for MRI; heart arrhythmia; glaucoma; hypertension; use of antidepressants or psychotropic medication; history of psychiatric illness or head trauma; drug or alcohol abuse; learning disabilities; poor eyesight; smoking more than 5 cigarettes a day; consumption of more than 24 units of alcohol per week; and pregnancy. All participants gave written informed consent before the experiment and were compensated with €110.

### Study design and MRI data acquisition

The study had a double-blind placebo-controlled crossover design. In each of two sessions, scheduled 1 week apart at the same time of day, participants received either a single oral dose of atomoxetine (40 mg) or placebo (microcrystalline cellulose PH 102, visually identical to the drug). Participants had a waiting time after pill ingestion, to allow for atomoxetine concentration to reach peak plasma levels. During the waiting interval, they either practiced the tasks (first session) or engaged in some quiet activity of their choice. Approximately 100 minutes after pill ingestion, participants underwent a resting-state scan, then performed one of the two tasks, underwent another resting-state scan, and performed the second task. The order of the two tasks was counterbalanced across participants.

All MRI data were collected with a Philips 3T MRI scanner. In the beginning of each of the scanning sessions, we collected a high-resolution anatomical T1 image (echo time 3.5 ms, repetition time 7.99 ms, flip angle 8°, and FOV 250 195 170 mm with voxel size of 1.1 mm isotropic). Functional scans consisted of T2*-weighted EPI images (echo time 30 ms, repetition time 2.2 s, flip angle 80°, FOV 220 220 120 mm with voxel size 2.75 mm isotropic). Eye tracking data were collected during the fMRI scans using an MRI-compatible EyeLink device. The camera was set to image the right eye and a calibration was carried out soon after the participant entered the scanner. The eye-tracking recordings were sampled at a frequency of 1000 Hz.

### Tasks and stimuli

In the scanner, participants underwent two resting state fMRI runs, and performed one run of a continuous performance task, and one run of a movie watching task. During the resting-state scan, participants were instructed to fixate their eyes on a black cross on a gray background. In the continuous performance task they were presented with a stream of images of cities and mountains, and had to mentally count the number of pictures with mountains. The stimuli consisted of 10 pictures of cities and 10 pictures of mountain landscapes that were presented multiple times in random order across the run. A key characteristic was that the transition between images was softened by making images fade into the next, which ensured a “continuous” type of stimulation. The images subtended approximately 6 degrees of visual angle, were isoluminant, grayscaled, and presented on a gray background. The images linearly and continuously morphed from one into the next, with an 800-ms interval (van den Brink et al. 2016). The percentage of mountains in the first session was set to 10% and on the second session to 40%. After the run, participants were asked how many mountains they had counted. The participants also performed a naturalistic viewing task which involved watching a shortened version of Hitchcock’s episode “Bang”. This clip has been widely used in fMRI experiments because it includes moments of suspense associated with specific brain patterns that can be observed across individuals (Naci et al. 2014). The clip included sound and subtitles. The resting-state runs and taskruns lasted approximately 8 minutes. The tasks were coded with Psychtoolbox running on MATLAB.

### Pupillometry preprocessing and analysis

Raw eye-tracking data were converted to asc files using the EDF2ASC converter. Preprocessing of the pupil signal consisted of interpolation of pupil blinks, normalization, detrending, and downsampling to the scanner frequency. Coding of these steps was based on code provided by Urai et al. (2017). Cross-correlation between pupil size and global signal was calculated with the function *xcorr* (MATLAB) as c = xcorr(global signal, pupil). We also identified *pseudo-events* in the resting-state pupil data, using the point process approach implemented in the RS-HRF toolbox (Wu et al. 2021). Briefly, the tool identifies the time points where the signal exceeds 1 standard deviation and fits an impulse response function. Thus, the *pupil events* are defined as pupil change that resemble an evoked response identified around points of high pupil signal. The RS-HRF toolbox allowed us to get the time points at which pupil events were detected. We calculated the proportion of pupil pseudo-events per time length of the intervals around global signal peaks (see below), because slower waves are expected to be related to longer intervals that may include more pupil events.

### fMRI preprocessing and analysis

Data were preprocessed using a standard pipeline implemented in FMRIPrep (Esteban et al. 2018, fmriprep.org), including slice-time correction, image realignment, deconfounding of movement regressors and mean signal of white matter and cerebrospinal fluid, detrending, and normalization to MNI space. Further steps included filtering (0.001-0.1 Hz) and z-scoring of the voxels’ timecourses. The analysis of brain activity propagation was done in surface space.

To study brain activity propagation along the principal gradient, we followed the procedure by Gu et al. (2021) and adapted the code shared by the authors. Briefly, the approach consists of, first, identifying the peaks of the global signal and the intervals around each of these peaks (i.e. from the trough before to the trough after each peak), and second, obtaining the time difference between peaks in the timecourses of the voxels with respect to the peak in the global signal, that is, the *delay values* (Figure 1). The vectors of delay values for all participants and scans were entered into a singular value decomposition algorithm. We then examined the first five main components (Figure 1) and identified the one corresponding to the *principal gradient*, that is, the one where brain regions are organized following a unimodal-transmodal gradient. Figure 1 shows the correlation between the vectors of delay values for all intervals considered and the vector corresponding to the principal gradient component. A bimodal distribution with a higher number of values in negative and positive correlations as compared to no correlation (zero value) indicates the presence of travelling waves in bottom-up and top-down directions along the gradient, respectively.

**Figure 1.**
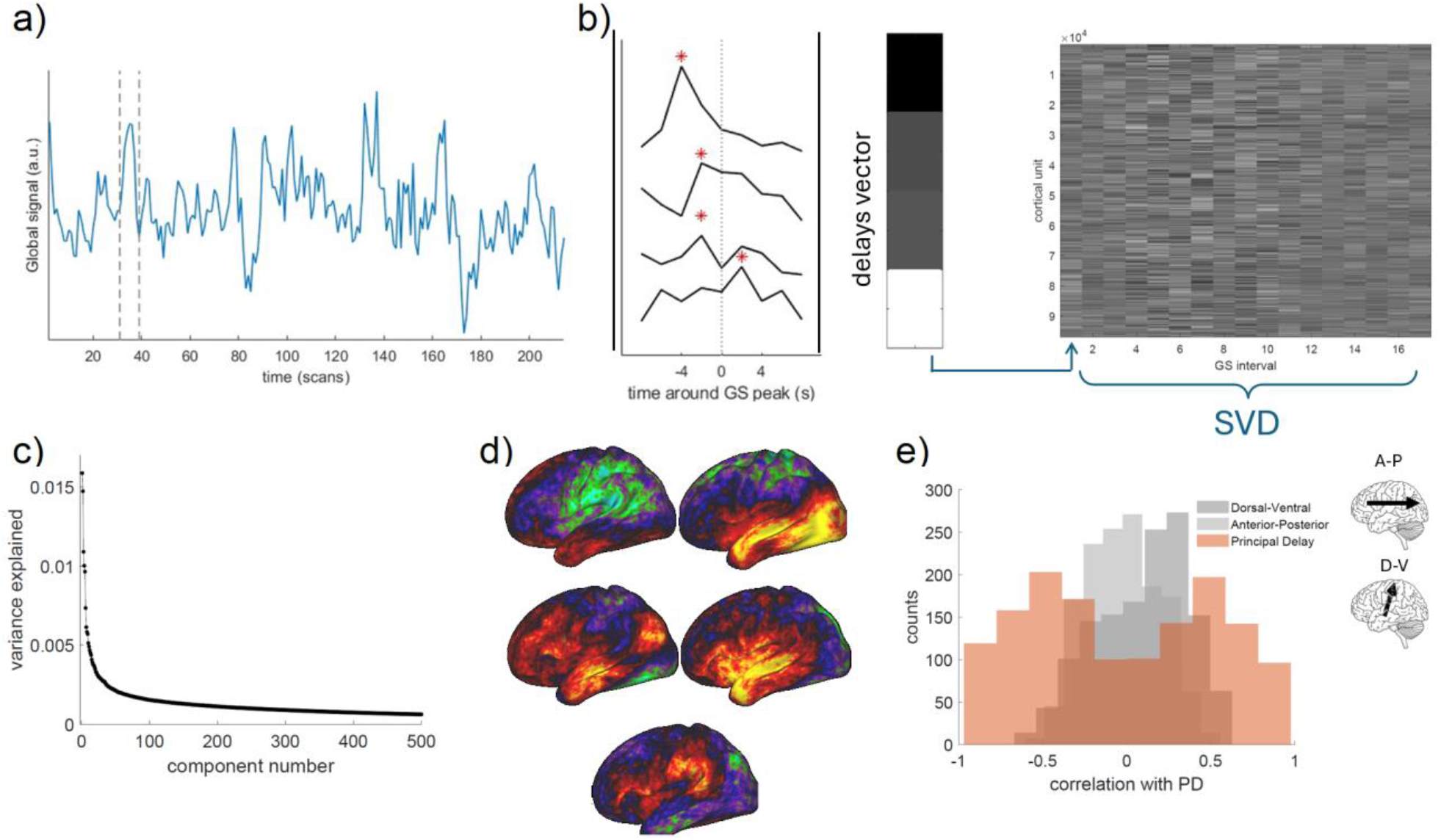
Calculation of main delay component. a) Example of extraction of delay vector from one session. b) The peaks of the global signal were identified (left) and the time difference between the peak of activity of each voxel and the global peak were calculated, resulting in one vector of delay values per interval (middle). The delay vectors were concatenated for all participants, sessions and tasks (right), and the matrix was submitted to a singular value decomposition analysis). c) Variance explained by each component. d) Brain maps corresponding to the first five components. The left map on the middle row (component #3) corresponds to the principal gradient. e) Histogram of correlation values between the delay vectors and the main delay component (red). The histograms corresponding to the anterior-posterior (A-P) and dorsal-ventral (D-V) gradients are also plotted. Brain insets indicate A-P and D-V directionality.

To study activity propagation along this gradient, for each individual and run, the voxels were ordered according to the gradient (i.e. from unimodal to multimodal) and binned in 30 bins to reduce the dimensionality of the analysis. The BOLD signal for each bin (i.e. averages over voxels) was used to study travelling waves (see a participant’s data example in Figure 1). Following Gu et al. (2021), we based the analyses on intervals of global involvement, for example those intervals where the global signal (averaged over all gray matter voxels) was larger than the peaks identified from a null distribution (i.e. based on shuffled data). To compute the speed of the waves, we used the formula as in Gu et al. (2021):

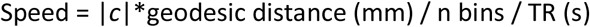

where the coefficient *c* corresponds to the linear fit where the position of the peak in the BOLD signal for each bin is regressed against bin number. A positive coefficient corresponded to a propagation from unimodal to transmodal areas (bottom-up), and a negative coefficient corresponded to a propagation from transmodal to unimodal (top-down). Figure 2 shows the average travelling waves in each direction. We set a threshold for the identification of travelling waves based on the distribution of coefficients for the two other propagation directions (A-P and V-D). The threshold was set at alpha = 0.1 and corresponded to a coefficient of absolute value 3.8 (Figure 2.d).

**Figure 2.**
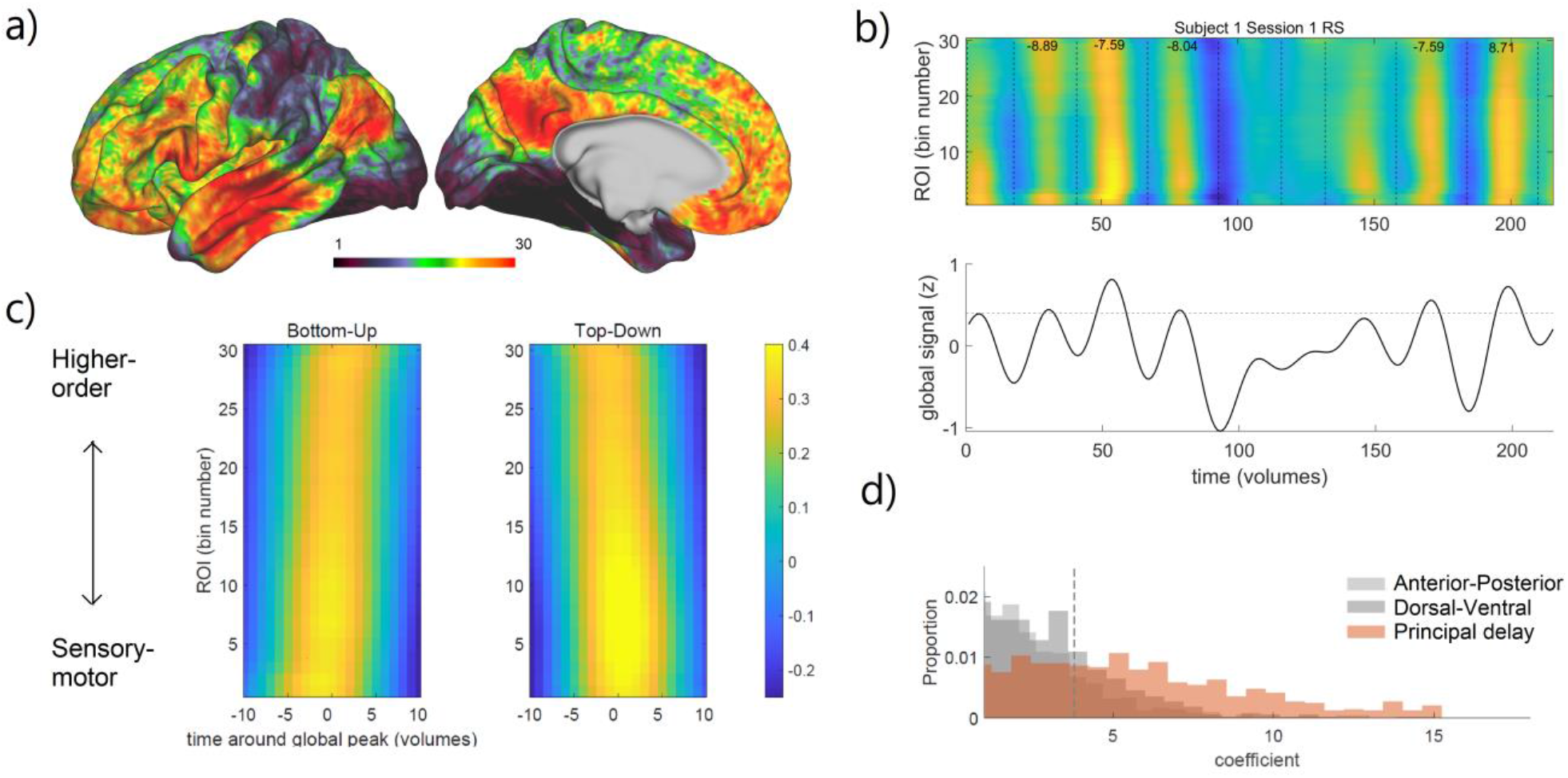
Calculation of speed of travelling waves. a) Map of principal delay component, binned into 30 bins according to delay with respect to the global signal peak. b) Example of one participant’s scan for which we plotted the BOLD signal of the regions of interest ordered after the principal delay (top panel). The travelling waves can be observed in relation to the global signal peaks (bottom panel). Horizontal line indicates significance threshold for global signal peaks obtained from permutation testing. c) Bottom-up and top-down travelling waves averaged across participants and conditions. d) Distribution of travelling wave coefficients for the two control directions and the principal delay direction. Threshold obtained from permutation testing is indicated with a vertical dashed line.

For the calculation of brain topological variables, we used the Gordon 333 parcellation (Gordon et al. 2016, Figure 6.a). In the Gordon 333 parcellation, each parcel is assigned to one of twelve different networks: Default, Parieto-Occipital, Fronto-Parietal, Salience, Cingulo-Opercular, Medial-Parietal, Dorsal-Attention, Ventral-Attention, Visual, Somatomotor hand, Somatomotor mouth, and Auditory. To account for the fact that the waves were related to peaks in global signal, which could inflate the estimation of integration measures, the global signal was regressed out from each voxel’s timecourse. Data from all voxels within a parcel were averaged. The temporal segments corresponding to the intervals with travelling waves were used to calculate the correlation coefficient between each pair of parcels, resulting in a connectivity matrix. The twelve networks were used as modules for the calculation of the participation coefficient, a measure of network integration. The participation coefficient was calculated as the average of the weights of a given parcel to each module with respect to all the weights of the parcel. The participation coefficient thus quantifies the extent to which a given parcel connects with parcels in other modules relative to parcels in its own module (van den Heuvel & Sporns 2013). A low participation coefficient corresponds to most of the connections of a parcel being distributed across a single or a small number of modules, while a high participation coefficient corresponds to connections of a parcel being distributed over a large proportion of modules. Therefore, the mean participation coefficient can be used to estimate the extent of integration of the entire brain network. For calculating the participation coefficient, we used the Brain Connectivity Toolbox (Rubinov and Sporns, 2010, https://sites.google.com/site/bctnet).

### Statistics

We were interested on the effects of treatment (atomoxetine vs. placebo) and behavioral state (rest vs. task-engaged) on travelling waves, and so we combined the data for the two resting-state scans and compared them with the aggregated data from the two tasks. We ran repeated-measures ANOVAs with propagation speed or top-down to bottom-up ratio as the dependent variable, and treatment and behavioral state as within-subject factors. For the analysis of propagation speed, we also included wave direction (top-down vs. bottom-up) as a within-subject factor. For the analysis of pupil-linked arousal, we ran a repeated-measures ANOVA with mean pupil size in a given time interval as the dependent variable, and treatment, behavioral state, and presence/direction of travelling waves (none, bottom-up, top-down) as within-subject factors. To test for a relation between propagation speed, mean pupil size, number of pseudo-pupil events (underlying the resting-state pupil signal), and measures of network-level integration, we ran a linear regression for each pair of variables, and included treatment and behavioral state as additional predictors. In these analyses we included all intervals, not only those of significant global involvement, so the correlations covered the whole range of variation in the pupil and integration variables. The linear models were coded using lme function in MATLAB.

## Results

### Effects of atomoxetine and behavioral state on brain activity propagation

First, we defined intervals around peaks in the global signal. Within those intervals, we then calculated the delay values of each voxel with respect to the global signal peak (delay profile). We finally applied singular value decomposition to those delay profiles to identify the main gradient of delays (Figure 1; Figure 3.c displays distribution of correlation values between the delay profiles and the principal gradient). For this, we used the whole dataset, including both treatments and behavioral states. Then, we binned the voxels into 30 equally-sized bins along the principal gradient and identified the bottom-up and top-down waves of activity propagation over the main delay profile in those intervals.

**Figure 3.**
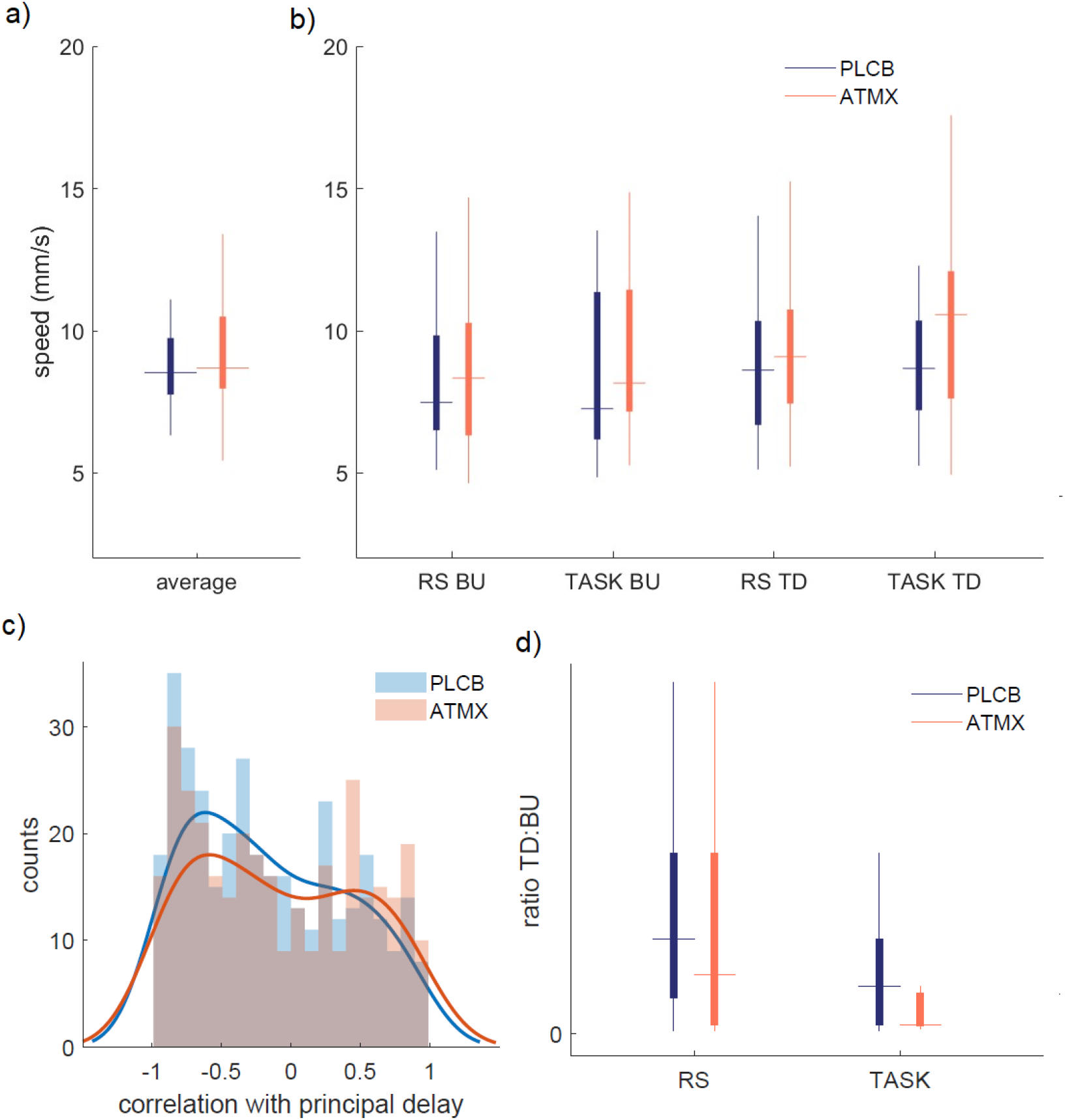
Travelling waves speed and ratio across treatment and behavioral state. a) Speed of travelling waves in the two different treatment conditions and b) separated by treatment, behavioral state and wave direction. c) Histogram of correlation values between the delay vectors of the individual windows with travelling waves and the principal delay vector. d) Ratio of top-down (TD) to bottom-up (BU) waves per treatment and behavioral state. In the boxplots, the central mark is the median, the edges of the box are the 25th and 75th percentiles, the whiskers extend to the most extreme datapoints. PLCB: placebo. ATMX: atomoxetine. RS: resting state.

To test our hypothesis that pharmacologically increased levels of catecholamines would be associated with faster travelling waves, we compared the propagation speed of bottom-up and top-down waves in the various conditions by performing a RM-ANOVA with the within-subject factors treatment, behavioral state and wave direction (bottom-up, BU, and top-down, TD). This analysis yielded a significant treatment effect (F(252) = 5.61, p = 0.018, Table 1). Speed was faster under atomoxetine as compared to placebo (Figure 3.a,b, Table 1), confirming our hypothesis. We found no effect of behavioral state or wave direction. Next, we compared the ratio of top-down to bottom-up travelling waves between conditions. This ratio significantly differed between behavioral states (F(110) = 10.50, p = 0.001), with smaller TD:BU ratio, that is, relatively more bottom-up waves, during task performance than rest (Figure 3.d). The ratio was numerically lower under atomoxetine as compared to placebo, but this effect was not significant (Table 1).

**Table 1.**

| Effect | Speed |  | TD:BU ratio |  |
| --- | --- | --- | --- | --- |
|  | F | P value | F | P value |
| Behavioral state | 1.28 | 0.257 | 10.54 | **0.001** |
| Treatment | 5.61 | **0.018** | 3.39 | 0.068 |
| Direction | 1.96 | 0.16 |  |  |
| Treatment* Behavioral state | 0.19 | 0.661 | 0.01 | 0.907 |
| Behavioral state*Direction | 0.18 | 0.670 |  |  |
| Treatment*Direction | 0.04 | 0.828 |  |  |

### Relationship between brain activity propagation and pupil-linked arousal

To study the relation between pupil size and brain dynamics, we investigated the slow fluctuations in pupil signal filtered with the same band-pass filter as the BOLD global signal (0.001-0.01 Hz). These fluctuations reflect slow changes in activity of neuromodulatory nuclei (Joshi & Gold, 2020; Mäki-Marttunen & Espeseth 2021; Lloyd et al., 2023, Figure 4.a). We found that the global signal and the filtered pupil signal were cross-correlated with a lag of around –5 scans (11 seconds), with fluctuations in the global signal preceding the corresponding fluctuations in pupil size (Figure 4.d). This direction is consistent with previous reports in mice (Pais-Roldán et al. 2020). The presence of several significant intervals may show certain periodicity in the signals. The cross-correlation was largely consistent across behavioral states and treatments. We also investigated pupil *pseudo-events*, that is, large (i.e., seemingly evoked) changes in the resting-state pupil signal (Figure 4.b, see Methods). We assumed that these were time points associated with phasic bursts of activity in neuromodulatory nuclei, including the locus coeruleus (Munn et al., 2021). The pseudo-events were only obtained in the resting-state scans to avoid stimulus-related pupil changes. We found that the pseudo-events clustered around global signal peaks, and in particular after the peak (Figure 4.c).

**Figure 4.**
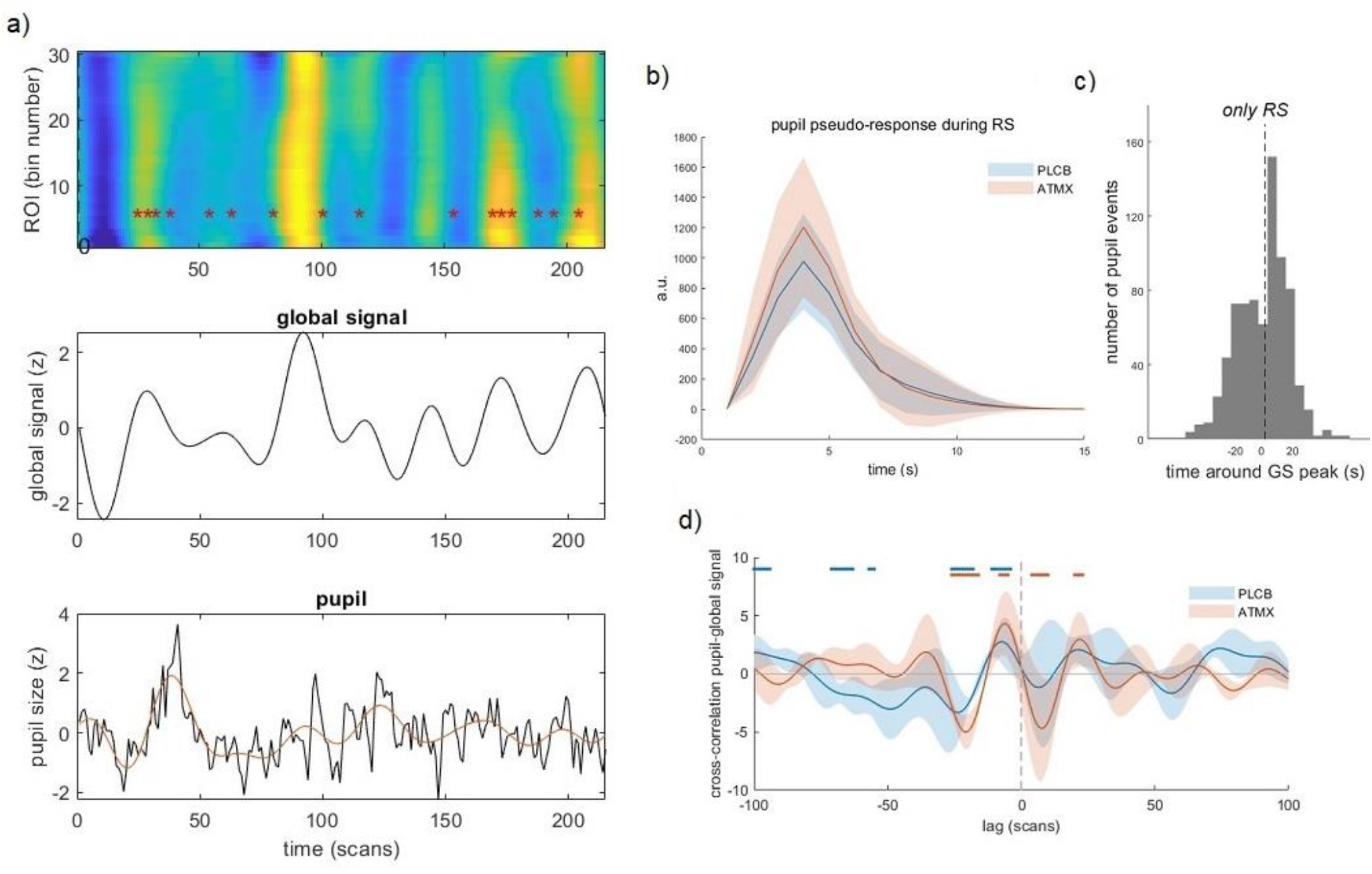
Relation between pupil size and global signal. a) A sample participant’s run. Top panel: brain activity ordered by the principal gradient with identified pupil pseudo-events (estimated from the resting-state pupil data) overlaid as red asterisks. Middle panel: global signal. Bottom panel: Pupil signal resampled to the fMRI sampling rate (black) and the low-passed pupil signal (red). b) Average of the phasic pupil changes that were identified as pseudo-events. c) Histogram of distribution of fast pupil events around global signal peaks. d) Average raw cross-correlation between pupil and global signal for the two treatments. PLCB: placebo. ATMX: atomoxetine. RS: resting state. GS: global signal.

To test our hypothesis that the presence of travelling waves would be associated with larger pupil size, we compared intervals with (bottom-up or top-down) and intervals without travelling waves. We found a significant effect, with larger pupil size in intervals with bottom-up or top-down travelling waves (Figure 5.a, presence of waves effect: F(277) = 4.19, p = 0.041), confirming our hypothesis. Note that the variability of mean pupil size was also larger for intervals with travelling waves. We then tested the relation between our pupil-related variables (mean pupil size and proportion of pseudo-events, Figure 5) and propagation speed. Propagation speed was not associated with mean pupil signal (F(1034) = 0.05, p = 0.822) but was significantly associated with proportion of pseudo-events (F(471) = 4.40, p = 0.036), whereby intervals with larger propagation speed were associated with less pseudo-events. No differences between conditions were observed in these analyses (mean pupil, treatment effect: F = 0.12, p = 0.723, behavioral state effect: F = 0.02, p = 0.877; pseudo-events, F = 0.049, p = 0.484). Taken together, the results suggest that time periods with traveling waves were characterized by larger mean pupil-linked arousal, and that speed of travelling waves was negatively associated with the proportion of pseudo-events.

**Figure 5.**
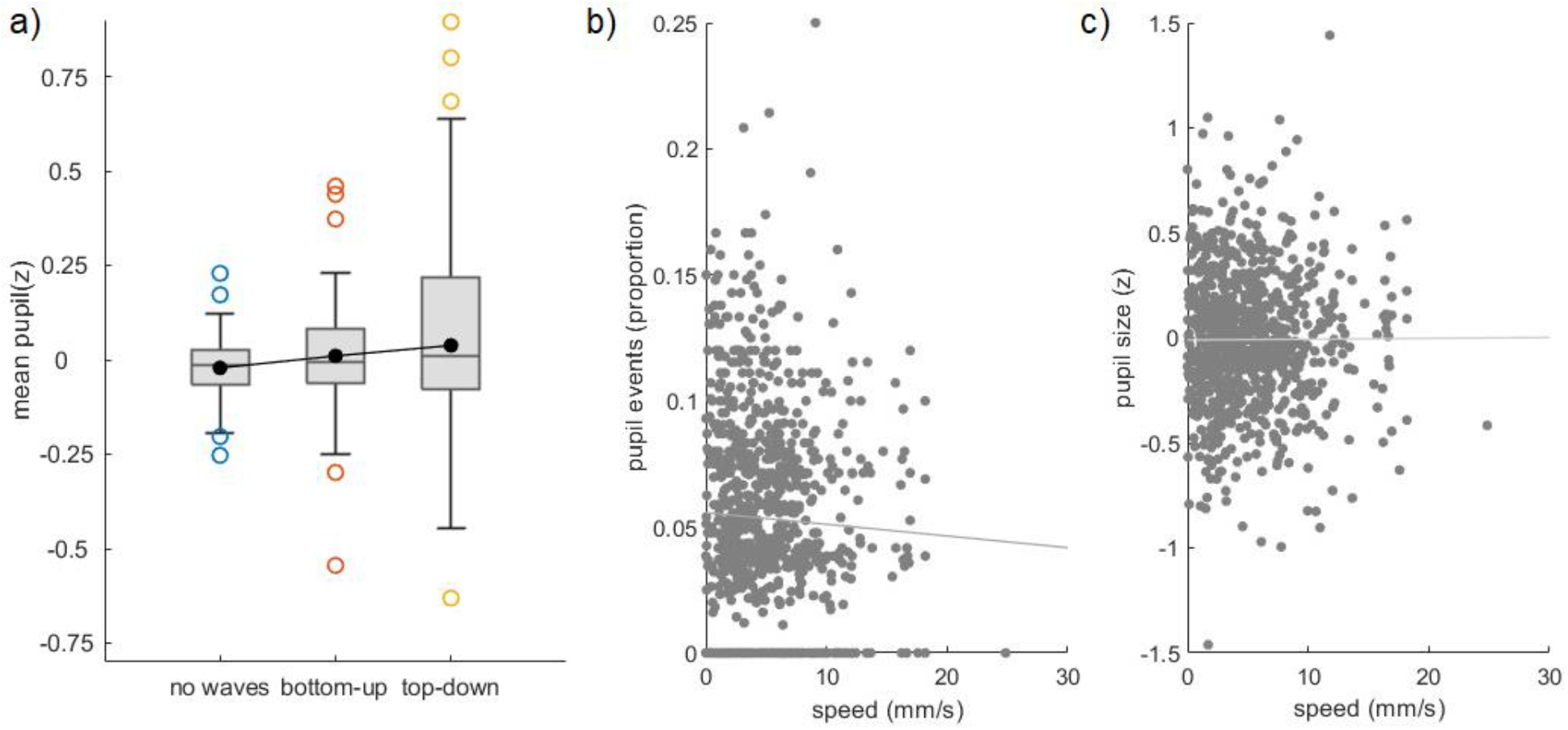
Links between travelling waves and pupil-linked arousal. a) Relation between presence/direction of travelling waves and mean pupil size. b) Relations between propagation speed and pupil measures. Each observation represents one interval.

**Figure 6.**
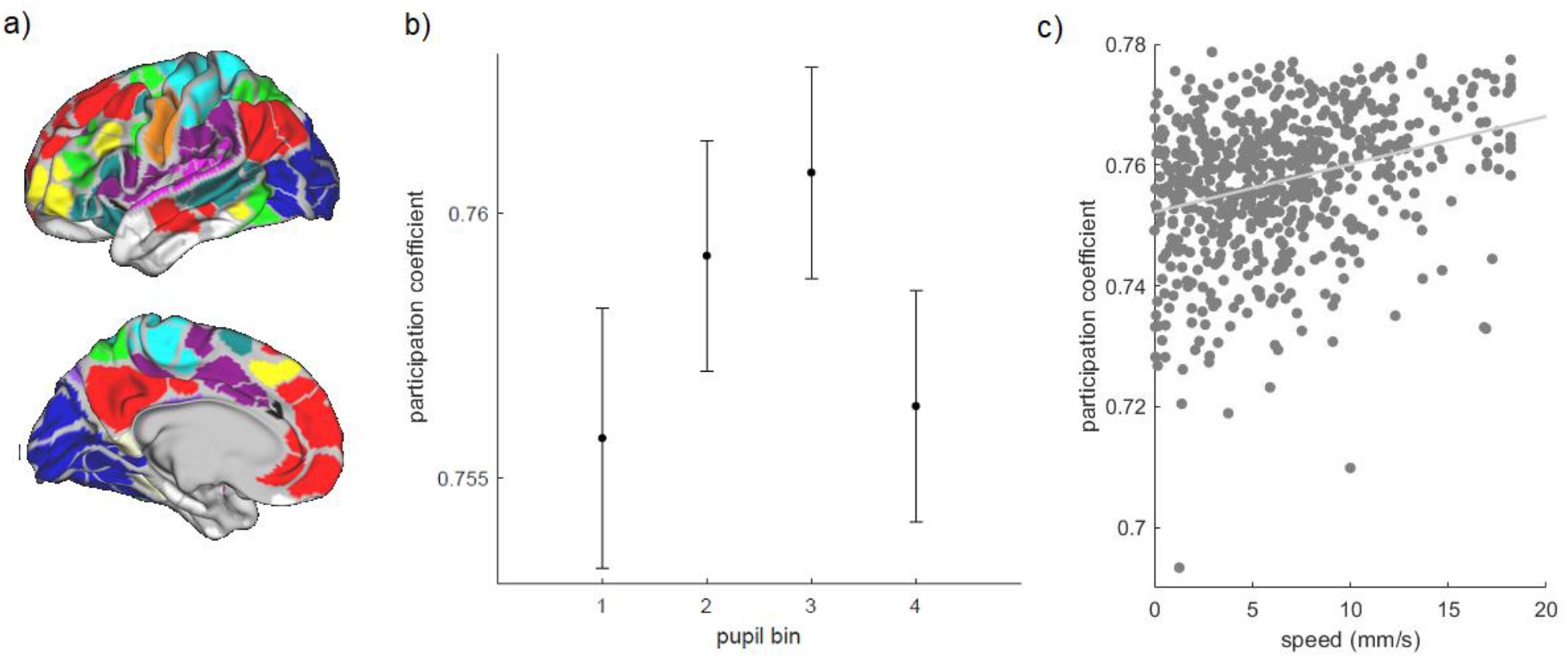
Brain integration. a) 333 parcells (Gordon et al. 2016) were used to calculate the participation coefficient, a measure of brain integration. b) Relation between pupil size and network integration. c) Relation between propagation speed and network integration.

### Relation between brain activity propagation and network topology

To test the hypothesis that that faster travelling wave speed would be associated with a more integrated network state, we calculated the participation coefficient, a measure of network integration. For this, we used the Gordon parcellation that divides up the cortex into 333 parcels (Figure 6.a) and averaged the data from all voxels within each parcel. Each parcel is assigned to a resting state network, and this assignment was used to compute the participation coefficient. We found that propagation speed was positively related to the level of integration (Figure 6.c; F(683) = 97.79, p < 0.001), confirming our hypothesis. There were no significant effects of treatment (F = 0.01, p = 0.925) and behavioral state (F = 0.74, p = 0.387) on level of integration. We also tested for a relation between pupil size and integration. We categorized all time intervals into four bins on the basis of mean pupil size and tested for an effect on the participation coefficient, following the procedure used in Mäki-Marttunen (2021). We found that larger pupil size was related to higher network integration, with the exception of the largest pupil bin, resulting in an inverted U shape (Figure 6.b, pupil bin linear effect: F(396) = 14.35, p < 0.001, pupil bin quadratic effect: F(396) = 13.83, p < 0.001). In sum, network integration was positively related to propagation speed and pupil size, the latter presenting also non-linear effects.

## Discussion

Propagation of brain activity in the form of travelling waves is a ubiquitous phenomenon of brain activity, although the factors that modulate them remain uncertain. The current study aimed to investigate whether neuromodulation has an enabling role on global activity propagation. To test this, our first approach was based on the proposal that the organization of brain activity into “networks” can be accounted for by the events of global propagation of activity (Raut et al. 2021), that the networks appear to fluctuate between moments of more integration across networks and moments of more segregation (Deco et al. 2015), and that pharmacological manipulations of neuromodulatory systems affect such measures of integration between functional networks (van den Brink et al. 2016, Shine et al. 2018). We had hypothesized that a **state** with higher levels of catecholamines would be associated with faster travelling waves, and that faster waves would lead to the measurement of more integration across different networks (Mäki-Marttunen 2023). Our second approach was based on the finding that moments of integration across functional networks are associated with moments of larger pupil size (Shine et al. 2016, Mäki-Marttunen 2021), which can be used as a **time-resolved** marker of slow changes in neuromodulatory activity. We hypothesized that this finding would be explained by larger pupil size being associated with the presence of travelling waves, with changes in arousal acting as an enabling mechanism of global propagation. Consistent with our hypotheses, we found that travelling waves were faster under atomoxetine, that faster travelling waves were related to more integration in functional connectivity, and that travelling waves were related to intervals of larger pupils. In sum, we found evidence in favor of the hypothesis that travelling waves underlie, at least in part, the organization of brain activity into functional connectivity networks, that the relationship between neuromodulatory levels and network integration is mediated by changes in the speed of propagation of travelling waves, and that moment-by-moment changes in pupil-linked arousal may enable or facilitate the emergence of global propagation of brain activity. This has important implications for the understanding of the brain mechanisms that underlie the dynamic properties of the human brain.

### Travelling waves

Here we examined the presence of travelling waves by extracting a main delay component of the BOLD signal based on intervals determined by global signal peaks and troughs, as in Gu et al. (2021). This method allowed us to obtain the main component that reproduces the main gradient of the brain, which separates unimodal from associative areas, and has been found across perceptual modalities and using various approaches (see Huntenburg et al. 2018 for a review). Propagation of activity in the form of travelling waves has been observed also in rodents and monkeys (Mitra et al. 2018, Matsui et al. 2016, Gu et al. 2021, Raut et al. 2021), indicating that they constitute a ubiquitous phenomenon. As in recent studies, we found both bottom-up and top-down propagation of activity. Importantly, we could detect them across the three behavioral states employed here: the resting state, the continuous performance task, and the movie watching. This is consistent with a previous study that found intrinsic patterns of activity following the principal gradient in resting state and a variety of tasks (Brown et al. 2022). An important difference is that in our study, none of the tasks required an active motor response or were paced as in event-related designs. Instead, they could be considered as inducing a “continuous” state, which made them comparable with the resting-state runs in terms of design. We found an effect of behavioral state on travelling wave ratio, where under both atomoxetine and placebo, tasks were associated with a larger proportion of bottom-up travelling waves than the resting state. Since the tasks had a strong visual component, it is possible that this effect depended in part on this property of the task (Brown et al. 2022). The atomoxetine manipulation was related to a larger proportion of bottom-up waves as compared to placebo, but this effect was non-significant (p = 0.068). A significantly larger proportion of bottom-up as compared to top-down travelling waves was found in sleepy individuals, as well as in monkeys (Gu et al. 2021). Recent studies using a serotoninergic agonist found that the principal gradient is contracted with increasing levels of serotonin (Girn et al. 2022, Timmermann et al., 2023). Taken together, the results suggest that neuromodulatory factors have an effect on the directionality of propagation of travelling waves along the principal gradient, reflecting the expression of a variety of brain states.

### Global brain activity, brain integration and neuromodulation

A key finding was that in both behavioral states, the speed of travelling waves was significantly higher in the atomoxetine condition. Computational studies suggest that one mechanism by which neuromodulatory systems influence the speed of travelling waves is by altering the overall excitability of the brain (Bhattacharya et al. 2021). The noradrenaline and dopamine systems, which are directly affected by noradrenaline transporter blockade with atomoxetine (Bymaster et al., 2002; Swanson et al., 2006; Koda et al., 2010), affect neuronal activity through binding to a variety of receptors distributed over the cortex (Aston-Jones and Waterhause, 2016; Avery and Krichmar, 2017; Rho et al. 2018). In particular, atomoxetine increases neuronal excitability (Bymaster et al., 2002; Koda et al., 2010; DiMiceli et al. 2015). Contrary to early views that noradrenaline affects all brain regions homogeneously, an interesting fact is that the receptors have different affinities and are distributed differentially over the cortex and the cortical layers (O’Donnell et al. 2012). In addition, projections from the noradrenergic nucleus locus coeruleus present some level of modularity, where locus coeruleus cells form ensembles that project each to specific regions (Chandler et al. 2019; Totah et al. 2018). Future work could disentangle whether the temporal modulation of travelling waves is related to the heterogeneity of noradrenergic projections. Overall, our results suggest that catecholaminergic activity exerts a nuanced modulation of brain activity propagation.

Studies using pupillometry concurrent with fMRI have found that periods of larger pupil size are related to more integration (Shine et al. 2016, Mäki-Marttunen 2021). Atomoxetine manipulation has also been found to affect the brain’s integration/segregation balance (Guedj et al. 2016, van den Brink et al. 2016, 2018, Shine et al. 2018). Different measures of brain integration were used across studies, which may underlie some differences in the direction of the effect. Our approach allowed us to look at neuromodulatory effects on a finer temporal scale by looking at the travelling waves over the principal gradient. The results revealed that periods of more integration may in part follow changes in speed of activity propagation across the principal gradient, and these may relate to fluctuations in neuromodulatory tone. Fluctuations between states of network integration and segregation have been described within single scan sessions (Betzel et al. 2016, Keilholz et al. 2013, Shine and Poldrack 2018). Furthermore, several studies reported that there are specific moments that contribute the most to the phenomenon of functional connectivity (Tagliazucchi et al. 2012; Liu and Duyn 2013). Our results allow tying together these lines of research by suggesting that the 1) modulation of macroscopic brain activity by neuromodulatory tone may reflect not only genuine changes in the overall state of functional connectivity but also the temporal relation of activity propagation across regions following the unimodal-transmodal hierarchy, 2) the effects of neuromodulatory activity on brain activity propagation may in turn underlie the fluctuations in functional network topology through modulation of propagating waves, and the single events of brain activity propagation in the form of travelling waves may be the ones particularly contributing to the emergence of brain functional connectivity. While future studies are needed to provide further support in favor of these hypotheses, our findings offer a clear picture of task-unrelated neuromodulatory effects on brain communication.

We also found that larger pupil size was related to more integration, although integration dropped at the largest pupil size values, following an inverted-U curve. Thus, at least within a certain range of pupil size, we replicated a previously reported positive relation between pupil size and netwrork integration, as quantified by the participation coefficient (Mäki-Marttunen 2021). We did not find an effect of pupil size on speed but found a significant negative association between number of pupil events and propagation speed. Pupil pseudo-events co-occur with BOLD changes in neuromodulatory regions in fMRI (Lloyd et al. 2023). While we can only speculate on the meaning of this finding, it is possible that time intervals of faster travelling waves are related to less variability in neuromodulatory activity and thus fewer pupil pseudo-events. While more work is needed to test for a relation between travelling waves and pupil size in other conditions and states, our results point towards a timed relation between neuromodulatory activity and global patterns of brain activity.

We also examined a possible relation between global signal and pupil size. The physiological meaning of global BOLD signal is still a matter of debate (Liu et al. 2017). The distinction between neuronal or artifactual sources of global signal changes has great relevance for the consideration of global signal removal as a preprocessing step in functional connectivity studies (Power et al. 2017). Previous studies suggest that the global signal is at least in part related to arousal level (Liu et al. 2018; Wong et al. 2013). We found a relation between pupil size, pupil pseudo-events and the global signal. A temporal relation between pupil size and global signal is consistent with previous reports in mice (Pais-Roldán et al. 2020). The fact that pupil size and pupil pseudo-events peaked after the global signal may reflect an event of transient arousal following a drop in arousal (Demiral et al. 2023). Our results contribute to the understanding of global brain activity by showing that pupil-linked fast and slow processes are related to fluctuations in BOLD signal. These pupil-linked processes may represent arousal influences at the peaks of global activity, that is, in a time-specific manner.

## Conclusions

By unveiling a relation between neuromodulatory tone, brain activity propagation along the unimodal-transmodal gradient, and pupil size, our study allows linking together several lines of research on brain dynamics, and provides a novel view of the effects of neruomodulatory systems across time scales.

